# Deciphering the Evolutionary Histories of Complex Genomic Rearrangements in Lung Malignancies

**DOI:** 10.1101/2024.11.11.623122

**Authors:** Xiaoju Hu, Vignesh Venkat, Gregory Riedlinger, Bing Xia, Missak Haigentz, Cristina Montagna, Zhiyuan Shen, Jyoti Malhotra, Subhajyoti De

**Affiliations:** Department of Pathology and Laboratory Medicine, Rutgers Cancer Institute, New Brunswick, NJ 08901, USA; Department of Radiation Oncology, Rutgers Cancer Institute, New Brunswick, NJ 08901, USA; Department of Medicine, Rutgers Cancer Institute, New Brunswick, NJ 08901, USA; Department of Medical Oncology & Therapeutics Research, City of Hope, Irvine, CA 92618, USA

## Abstract

Complex genomic rearrangements (CGRs) are hallmarks of tumor genomes, yet their architectures and origins remain poorly understood. We utilized optical mapping to profile the genomic landscapes of lung tissues across various stages - from normal and benign lesions to advanced carcinomas - to characterize genome disorganization during carcinogenesis. We developed omcplR to resolve the CGR architectures and predict their genesis using the genome-graph framework. Our findings reveal that many CGRs arise from hierarchical combinations of localized structural variations that are unlikely due to break-fusion-bridge, chromothripsis, or chromoplexy. We termed this process chromodiadochi: a cascade of nested rearrangements accumulating over multiple generations during carcinogenesis. Early intra-chromosomal junctions in CGRs typically localized within specific chromatin domains, while late junctions in advanced tumors occurred in more heterogeneous contexts and showed signs of positive selection. A composite signature of genomic alterations, including the CGRs, captures the trajectory of progressive genomic structural disorganization and instability from benign tissue to malignancy.

## Introduction

Genomic instability and mutagenesis are hallmarks of cancer, and most tumors have rearranged genomes^1–4^. While more than half of somatic point mutations and small InDels detected in cancer genomes are suspected to arise prior to malignancy during development and aging^5^, genomic rearrangements show very different patterns – they are rare in normal tissues but prevalent in tumors. Most genomic rearrangements in tumors are structurally simple, but complex genomic rearrangements (CGR) i.e. genomic reorganization events involving multiple intertwined structural rearrangements are common in all major cancer types^4^. Recent pan-cancer studies, primarily based on short read sequencing, have identified different mechanisms of complex rearrangements such as chromothripsis, break-bridge-fusion, chromoplexy, etc, which have been collectively termed as chromoanagenesis^2,6–8^. The functional significance of CGRs in the context of carcinogenesis appears to be high. Focal amplifications of many known oncogenic drivers (e.g., MET, EGFR, ERBB2) frequently involve complex rearrangements, and their allelic heterogeneity provides substrates for adaptive evolution and acquired therapeutic resistance^4,9^. A subset of the complex rearrangements may be associated with micronuclei or extrachromosomal circularized DNA copies; these structures are subject to unequal segregation during cell division, allowing rapid and reversible gains or losses of copy number that can confer dynamic intra-tumor heterogeneity, facilitating rapid tumor evolution, and adaptive treatment resistance^10^.

Despite their clinical significance, the genomic architectures and molecular mechanisms behind the CGRs are poorly understood. Traditional DNA FISH can detect translocations and copy number alterations, but lacks the resolution to resolve the structure of complex rearrangements. Short read sequencing is routinely used for profiling tumor genomes; however, it has been debated^10,11^ whether this technology can effectively identify rearrangement junctions in repetitive regions or the connected genomic alteration events occurring on the same allelic haplotype, especially if such events are kb-Mb apart. Long read technology allows sequencing long DNA fragments, but efforts to reconstruct the architecture of complex genomic arrangements have been lagging. To address this gap, we took advantage of ultra-long read DNA profiling using ultra-long optical mapping (OM) technology and developed omcplR to resolve the architectures of CGRs.

Unusual complexity of the genomic architecture of the CGRs raises several fundamental questions: when and how do these events arise during tumorigenesis? Are these events typically associated with genomic instability and remain under selection? Answers to these questions require examining the genomic landscapes of premalignant and malignant lesions to assess the complexity of the CGRs, considering progressive loss of genomic organization and onset of genome instability during malignancy^12,13^. Unfortunately, early tumor development is typically asymptomatic, not all benign growths progress to advanced disease stages, and the biobanking of relevant premalignant lesions is typically limited. Lung malignancies are notable exceptions; pulmonary nodules are often detected during lung cancer screening, but only a subset of the pulmonary nodules of indeterminate potential progress to become malignant (**Figure 1A**), even though all benign tumors progress to become cancerous ^14,15^; classifying the nodules with indeterminate potential remains challenging. We used OM to profile the genomes of tissue samples from different stages of lung malignancies to identify the CGRs and then apply omcplR to examine their etiologies and evolutionary histories.

**Figure 1:**
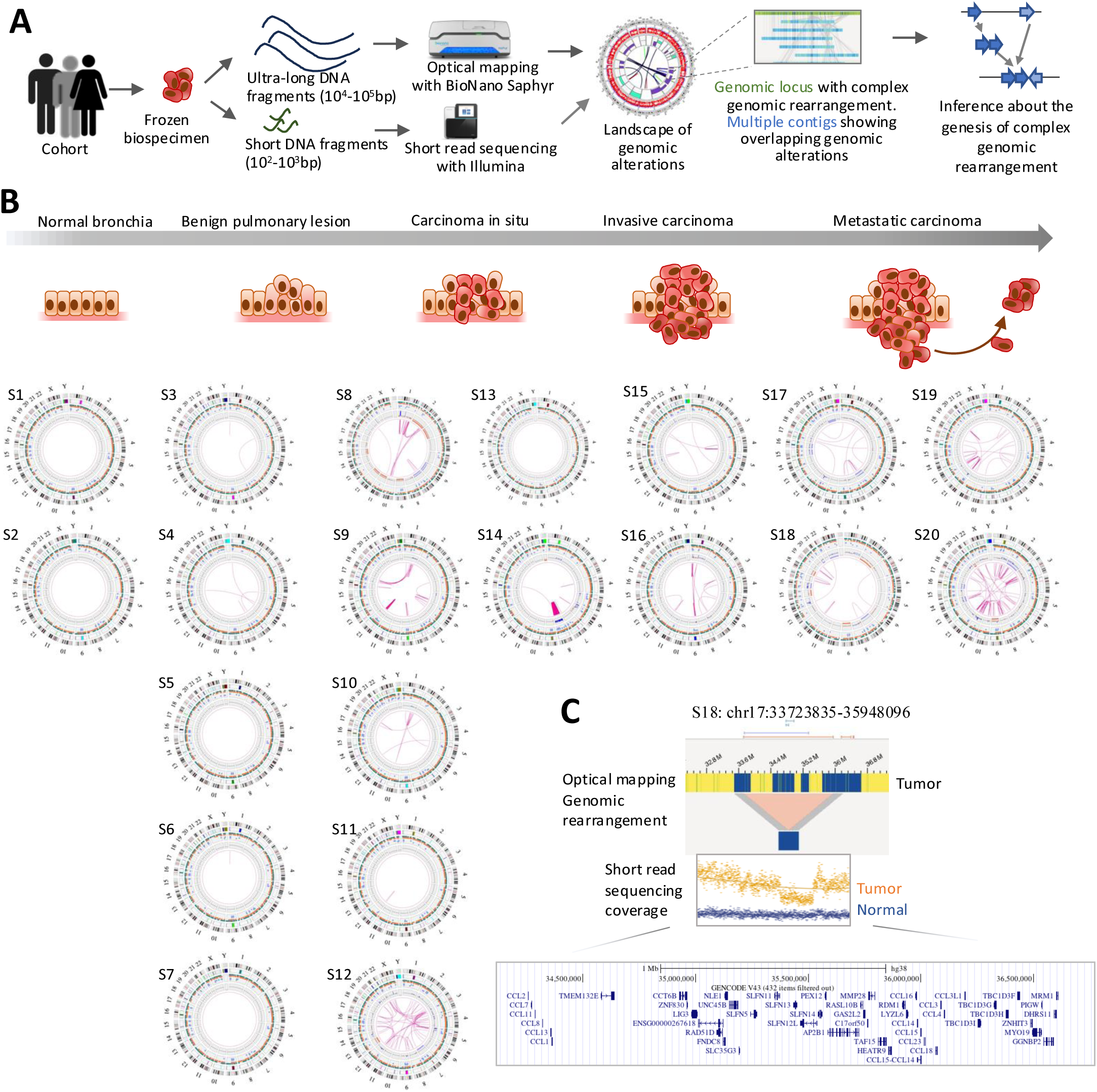
Profiling of the landscape of genomic rearrangements in lung tissues using optical mapping. A) Outline of the study design profiling genomic landscapes of pathologically normal lung tissues, benign dysplastic and hyperplastic lesions, and malignant lung carcinoma samples. B) Circos plots showing somat ic genomic structural alterations and rearrangements in the samples in the cohort, grouped according to the disease stages. C) Representative genomic alteration in a sample (S18) is shown at high resolution, and corresponding variation in sequencing coverage in Illumina short read sequencing of the tumor is highlighted (tumor: orange, paired normal: blue). The genomic resolution of optical mapping and short read sequencing are different. The affected and adjoining genomic regions and gene annotations therein are shown below.

## Results

### Cohort characteristics and pathobiology of the samples

We collected well-curated, fresh frozen tissue samples from pathologically normal lung tissues, benign pulmonary lesions, carcinoma in situ, locally invasive carcinomas, and metastatic lung carcinomas. These samples were obtained from a cohort of deidentified donors (**Figure 1B**, **Supplementary Table 1**) under an IRB-approved protocol. Lung tumors and adjacent pathologically normal tissues were sourced from cancer patients, while benign pulmonary lesions were collected during preventive surgeries.

The benign lesions were typically identified as suspicious nodules of indeterminate potential and exhibited additional characteristics associated with elevated cancer risk. For instance, Sample S5 was a 1.5 x 1.3 x 1.2 cm mass with ill-defined boundaries and a firm consistency. It showed diffuse interstitial involvement by non-necrotizing granulomatous inflammation, for which stains for infectious agents were negative. Sample S6 was a 3.5 x 2 x 1.1 cm hard, calcified nodule with focal atypical cells, suggestive of abnormal pulmonary growth. This patient had a prior history of squamous cell carcinoma and had received chemoradiation; the malignant tumor was removed via wedge resection and did not overlap with the nodule. Sample S4 was a well-circumscribed 0.5 x 0.5 x 0.4 cm non-malignant pulmonary nodule collected via wedge resection. Regional inflammation was noted for several benign lesions, but stains for infectious agents were consistently negative.

Among the cancerous lesions, approximately one-third presented with locally invasive and/or metastatic disease, while the remainder were classified as carcinoma in situ. Histological characteristics indicated the presence of papillary, lepidic, and solid subtypes, with some tumors exhibiting mixed histology. For example, Sample S9 was a 2.0 x 1.7 x 1.9 cm infiltrating adenocarcinoma of the lung with lepidic (50%), papillary (5%), and acinar (45%) patterns; no lymph node or pleural invasion was detected. This patient had a prior history of prostate cancer. Sample S17 was diagnosed as a 1.6 cm invasive mucin-producing adenocarcinoma with mixed subtypes, including lepidic, acinar, and papillary patterns, associated with granulomatous inflammation; invasion of the visceral pleura was noted. Sample S20 was an invasive, moderately differentiated lung adenocarcinoma, predominantly acinar, with lymph node metastasis (pT1cN1).

### OM-based genomic profiling of the cohort

We performed optical genome mapping at ∼350X effective coverage on the frozen samples using the BioNano Saphyr platform^16^. Bionano Optical Genome Mapping directly observes genomic regions and structural variations by linearizing and imaging DNA in its native state using massively parallel nano-channels. OM provides a detailed "map" of individual single DNA molecules by observing the pattern of fluorescent tags along their lengths, potentially providing high-resolution information about structural variations. The optical mapping (OM) data were processed and analyzed after multiple QC filters (**Supplementary Table 2**, see **Methods** for details). On average, the total length of the input maps was 2,375,219 ±526,469.2 Mb, with an N50 length of 205.35±47.30 kb, and more than 91.55% ± 6.52% of the genome maps could be mapped to the human reference genome (GRCh38), achieving a genome coverage of 351.76±45.90X. Some genomic regions had multiple contigs representing multiple alternate alleles due to the presence of one or more genomic alteration events. We applied the BNG rare variant analysis pipeline to identify different classes of structural variations and copy number alterations. The landscapes of genomic rearrangements and other alterations in these samples are presented using circos plots (**Figure 1B**). The insertions (median: 5.45kb) and deletions (median: 16.79kb) were the most prevalent variants reported in the dataset, while intra-chromosomal fusions and inter-chromosomal translocations were less common in the cohort, and more so in the non-malignant samples.

For comparative assessment and validation, we performed whole genome sequencing (WGS) of 4 samples -2 sets of advanced stage carcinomas (S8 and S18) and paired normal tissues (S1 and S2) using 150bp short read PE sequencing on the Illumina platform (see Methods for details). The number of detectable somatic point mutations (S8: 33,541, S18: 18,445) and InDels (S8: 1396, S18: 570) were within the typical range expected for lung carcinomas^17^. WGS using short read technology identified several structural variations in the tumor samples (n=40 for S8, and n=15 for S18; **Supplementary Table 3**), but the frequency was lower than that identified using the OM technology. Furthermore, a vast majority of the high-confidence somatic structural variations in tumors identified on the short-read platform were below the size threshold for detection using the OM technology. Among the large structural variations within the detection range of both technologies, those called by WGS were also detected by OM. For instance, S18 has one deletion in chr17: 33.7Mb-35.9Mb that was found in both platforms (**Figure 1C**). We identified four SVs (two inversions and two deletions), which were detected by both platforms (**Supplementary Table 4**). In contrast, most of the high-confidence, high allele frequency (>10%) OM structural variation calls were not identified by WGS. In some cases, genomic evidence on the Illumina platform was below the confidence level for detection threshold, and higher depth of coverage could be necessary (also see **Discussion**), while some others had rearrangement junctions in the repetitive or low complexity sequences that are difficult to resolve using short read technologies. Nonetheless, our observations about the extent of concordance between the platforms are similar to those reported in other studies^18,19^.

A comparative analysis of the genomic landscapes of the samples profiled using OM across different disease stages (**Figure 1B**) indicated that large copy number alterations and inter-chromosomal rearrangements were rare in pathologically normal tissues and benign samples, but prevalent in the in-situ and advanced carcinomas. This is consistent with the observations on the mutational landscapes of healthy, diseased, and malignant lung samples profiled using short read technologies^17,20^. While a significant burden of somatic point mutations and small InDels in the genomes of lung cancer, and that of most solid tumors, arises prior to malignancy during development and aging^5,12,15^, these observations collectively suggest that large genomic rearrangements are typically rare in non-malignant tissues, and are characteristic of malignant disease progression.

The contigs based on the long OM molecular traces allowed detection of both junctions of the rearrangements, even if those are 10^4^-10^6^bp apart, enabling end-to-end characterization and reconstruction of the architecture of the genomic alteration events. Based on the contig-level data, we further identified the instances where multiple rearrangement and/or copy number alteration events overlap or occur within the same OM contig and tagged those as clustered SVs. We identified 25-36 clustered SVs in the benign lesions, 24-59 in carcinoma in situ, and 32-59 in the invasive tumors and those with metastasis (**Supplementary Table 5**), respectively. Most clustered SVs consisted of relatively small, localized structural variations, including deletions, insertions, duplications, and inversions, but some of the clusters spanned local (∼hundreds of kb) or distal (∼tens of Mb) regions of chromosomes, and/or involved inter-chromosomal rearrangement events. An analysis of the junction points and visualization on the Bionano VIA platform indicated that these clustered structural variations (SVs) likely represent complex genomic rearrangements, resulting from multiple entangled alteration events on a single allelic contig.

We then annotated known cancer genes proximal to genomic rearrangement junctions. Clustered rearrangements involving U2AF1, IRF4, ZNF521, PDE4DIP, HIP1, WWTR1, NOTCH2, NRG1, and MAPK1 were identified across multiple disease stages. While U2AF1- and HIP1-proximal events were frequently observed in invasive and distant carcinomas (**Supplementary Table 6**), this enrichment did not reach statistical significance. Given the specific resolution of OM and the absence of point mutation data, this gene set likely represents a subset of the total oncogenic drivers in this cohort.

Although focal amplifications of MET and ERBB2 - common in non-small cell lung carcinoma (NSCLC)^17^ - were not detected via OM, single-cell transcriptomic data for these samples^21^ revealed significant oncogenic pathway activities. Advanced carcinomas S17 and S20 exhibited elevated EGFR pathway expression, suggesting potential RTK signaling activation; notably, high EGFR expression was also present in the benign lesion S7. Furthermore, samples S19 (acinar), S11, and S10 (papillary) demonstrated high KRAS expression, consistent with the high frequency of KRAS mutations typically associated with these histological subtypes^22^.

### omcplR for characterization of complex genomic rearrangements

We developed omcplR to identify and resolve CGR architectures from optical mapping (OM) data (see **Methods**). Some complex rearrangements may form in a single mutagenic event, while others may develop due to recurrent DNA double-strand breaks leading to overlapping rearrangements at a locus over extended period during tumorigenesis. To distinguish between different temporal patterns, omcpIR analyzed the rearrangement architecture within the CGRs by considering the relationships among the rearrangement junctions therein.

OmcplR first partitioned the genome into equal-sized segments to identify "seeds"—regions where multiple contigs harbor divergent structural variation (SV) junctions. Using a recursive search, omcplR iteratively clustered co-occurring contigs until no new member could be added to a cluster. We then extracted SV junctions from these clusters and retained unique contigs as alternative alleles. Each resulting non-redundant set defines a distinct complex genomic rearrangement (CGR). Within a CGR, some of the contigs carried multiple rearrangement junctions, and on the other hand, a subset of the rearrangement junctions was shared across multiple contigs. omcpIR used the joint patterns of contigs and rearrangement junctions to infer the conditional relationships among the junctions, and thereby predict the likely order of precedence of rearrangement events therein (**Figure 2A**). For each CGR set, omcpIR estimated the frequency of pairwise co-occurrence between the junctions, and then used a Bayesian approach to determine conditional relationships among all possible junction pairs – such that the CGR set was graphically represented as a Bayesian network, where the nodes were SV junctions and the pairs of junctions with significant conditional relationships related to edges (see Methods for details). In brief, a complete undirected graph, where every node is connected to every other node, was constructed, and then the edges were systematically removed based on a series of conditional independence tests. Finally, as many of the remaining undirected edges were converted into directed edges as possible based on the conditional co-occurrence without creating cycles in the graph to generate a Completed Partially Directed Acyclic Graph (CPDAG). The graph represents the conditional relationships among the rearrangement events within a CGR.

**Figure 2:**
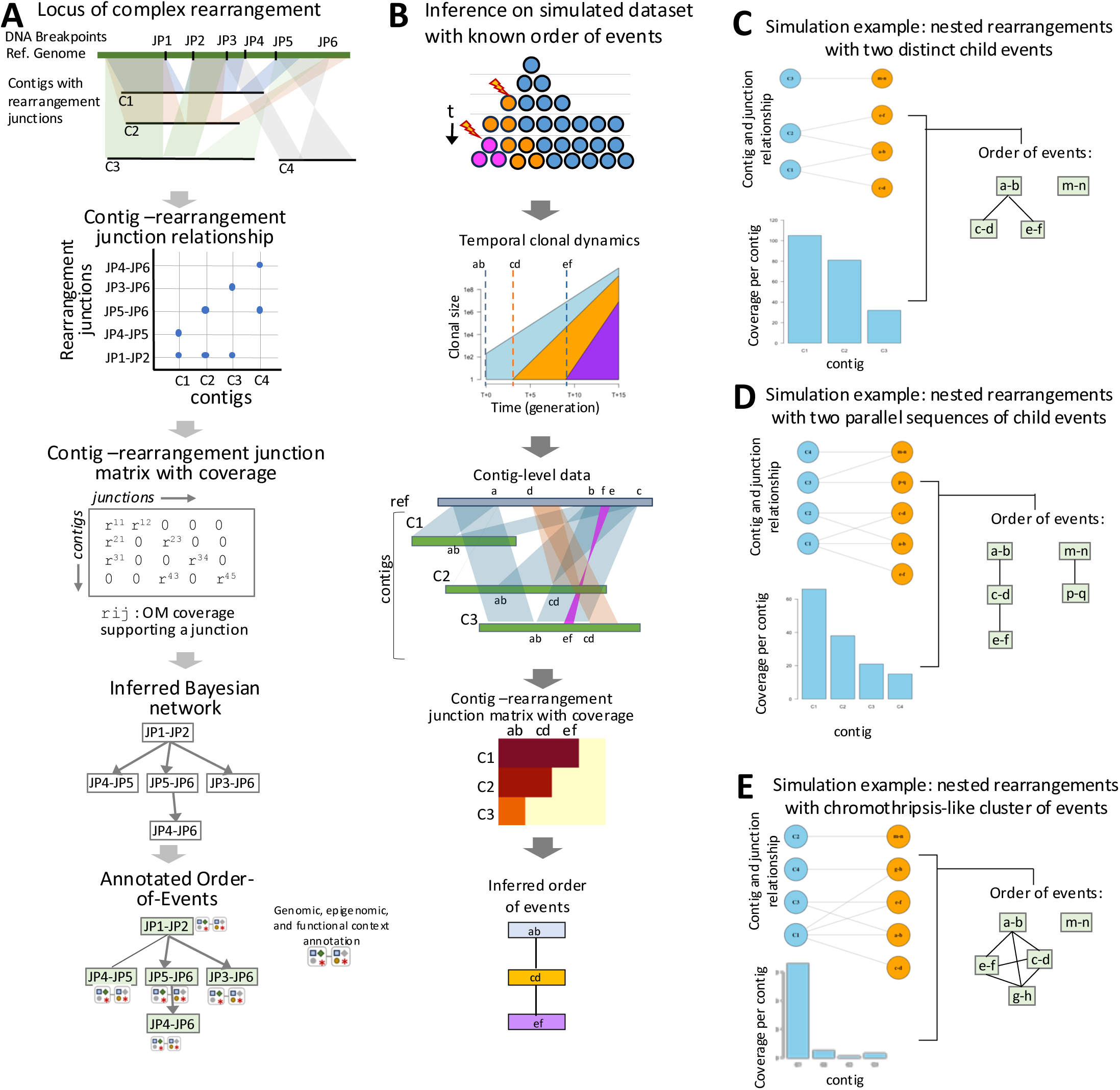
Order-of-events inference using omcplR framework. A) A schematic representation of the computational pipeline to identify complex genomic rearrangements and infer underlying order of events. B) Benchmarking using simulated datasets where the order of rearrangement events is known in advance. C-E) Order-of-inference based on simulation, where nested rearrangements with C) two distinct child events, D) two parallel sequences of child events, and chromothripsis-like cluster of events were predicted. The rearrangement junctions are shown in orange, while the contigs are shown in blue. The edges show their respective relationships.

The likely sequence of rearrangement events during tumorigenesis could be inferred from the topology of the network top-down, considering the conditional dependence of the junction-points – early rearrangement junctions were at or near the top of the graphs, while downstream events were below, such that one may annotate the rearrangement junctions within a CGR hierarchy as early, intermediate, or late (**Figure 2A**). Outside the direct decedent relationships, the order-of-events inference needs cautious interpretation. If a complex event has multiple disjoint graphs, no statistical inference can be made regarding the relative temporal order of occurrence of rearrangement events across the graphs. Likewise, no inference could be made regarding temporal order between different unrelated CGRs in a cancer genome.

We first evaluated the performance of omcpIR using computational simulation, where the ground truth about the order of events was known in advance (see **Methods**). While a single catastrophic CGR event would have comparable allelic counts of the interconnected events, those that arose due to temporally distinct events would have different allelic counts within the same CGR. Using different model parameters, representing different evolutionary scenarios and noise-level in the OM data, we propagated the clonal lineages in the simulation, keeping track of the CGRs, their order of events, and tabulated the final allelic counts of the genomic alterations therein (**Figure 2B**). We found that omcpIR could correctly distinguish those that likely arose via sequential accumulation of rearrangements and identify the order of events in complex CGRs, and present three examples (**Figure 2C-E**). The first example has nested rearrangements with two distinct child events after an early event, as evident from their contig-level membership; this was correctly established by omcplR (**Figure 2C**). Likewise, we simulated a complex locus, which had two parallel sequences of child events (**Figure 2D**), and the algorithm predicted their probable sequence of events correctly. Finally, we simulated a chromothripsis-like cluster of events (**Figure 2E**), and indeed, omcplR reported no order of precedence. In this analysis, the addition of a low level of technical noise, in the form of variations in the local coverage or the presence of low coverage contigs with only a subset of the junction-points - did not alter the conclusions. We evaluated the model with different constraints, coverage, and noise levels (**Supplementary Table 7**). The model demonstrated high fidelity in reconstructing temporal sequences when analyzing event-level count data, especially the event sequence is well-defined and the events are sufficiently separated along the clonal lineages.

However, inference uncertainty can be impacted by local coverage, dropouts in the structural variant calls, and noise in the count-level data, such that distinguishing singular catastrophic events from those that arose via successive events during carcinogenesis may be challenging.

Next, we scanned the somatic genomes of the premalignant lesions, as well as localized and advanced carcinomas in our cohort, using omcpIR to identify the CGRs and predict their likely order of events. Most events were heterologous, i.e., involving multiple classes of genomic alterations and involved multiple overlapping contigs – which reflected diversity and heterogeneity within the CGR. A majority of the CGRs were intra-chromosomal localized within 1Mb regions, such that the rearrangements were within the same topological domains and might not perturb long-range functional interactions within the rearranged regions. Distal (>1Mb) intra-chromosomal events were more common in malignant tumors. As we discuss later, the burden of inter-chromosomal complex events increased from non-malignant to malignant samples.

As a case study, we showcase a complex genomic rearrangement event located in chromosome 15 between 23.13Mb-28.62Mb of distant tumor S20, which is an advanced carcinoma (**Figure 3A**; see **Methods** for details). This CGR involved 7 alternative DNA contigs that harbored different combinations of structural variations, often involving common subsets of rearrangement junctions (**Supplementary Table 8**). We show the joint distributions of the rearrangement junctions on the contigs using a circus plot, where the width of the arcs indicates the extent of co-occurrence of rearrangement junction-pairs in that sample, involving the most prevalent joint junctions: Chr15_28.42MB||Chr15_28.42MB - Chr15_28.50MB||Chr15_28.62MB (**Supplementary Table 8**). Gene-level annotation suggested that the CGR impacted the genes GOLGA8DP, GOLGA6L2, GOLGA6L7, and MKRN3. The rearrangement junctions in this CGR were within the same chromatin domain and shared similar (epi)genomic contexts (**Figure 3B**). All the junctions were in the strong euchromatin context in the nuclear core, and nearly all of them overlapped with repetitive elements (2 junctions lacked nuclear context annotations, Chr15:23.13MB||Chr15:23.13MB and Chr15:23.36MB||Chr15:28.34MB). Close sequence-level and topological proximity of all involved rearrangement junctions are consistent with a model that DNA breaks within a replication factory and recursive rearrangements could give rise to such events (**Figure 3C**). Furthermore, the order of events analysis of this complex region indicated that the segment junctions Chr15:23.13MB||Chr15:23.13MB and Chr15:23.59MB||Chr15:28.51MB might be the early event (**Figure 3B**). Close sequence-level and topological proximity of all involved rearrangement junctions within the same chromatin domain context suggest that the DNA breaks likely occurred within a common replication factory, and we speculate that recursive DNA breaks perhaps due to persistent replication factory crisis^23^ might lead to such events (**Figure 3C**).

**Figure 3:**
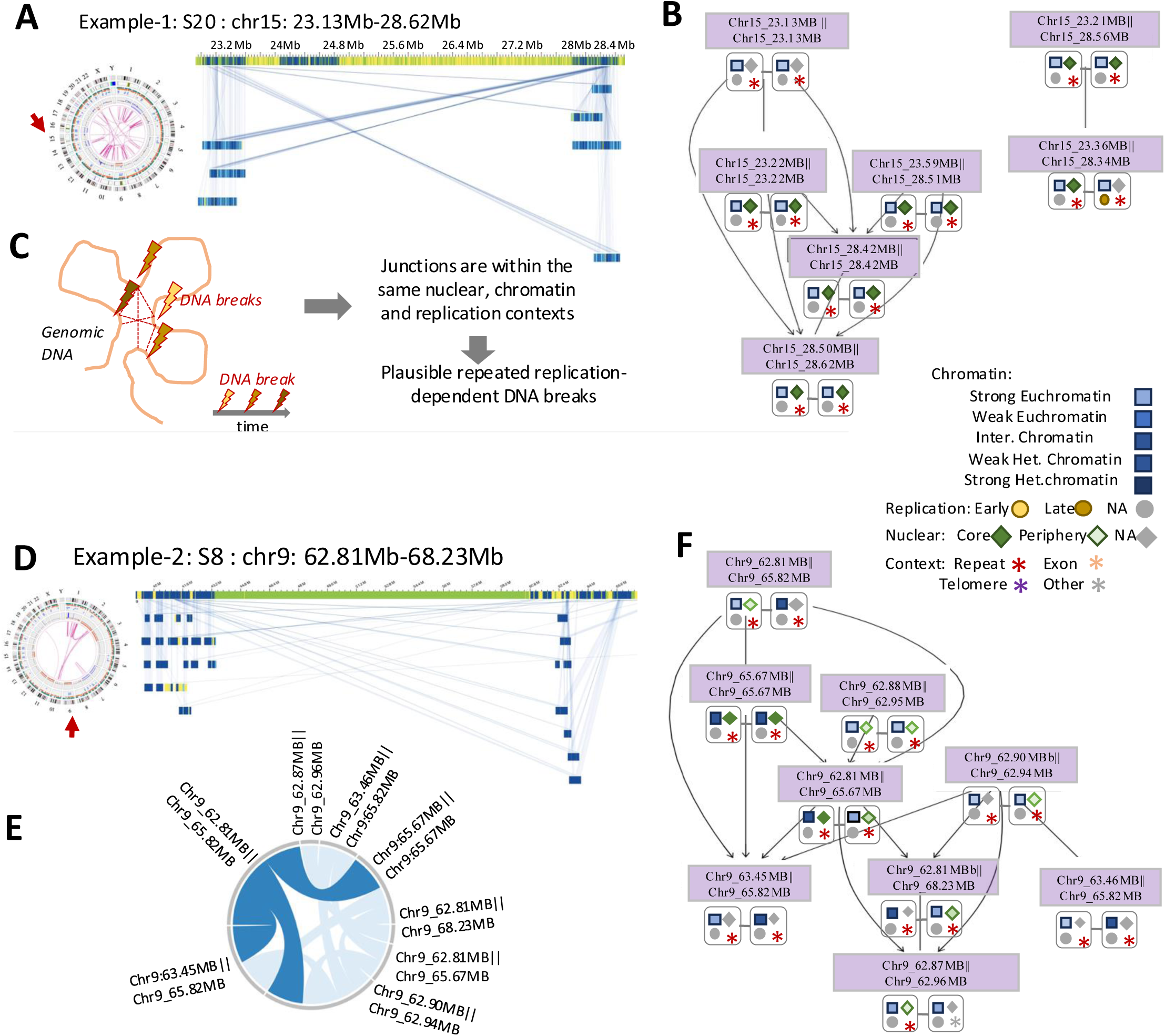
Genomic architecture and order-of-events inference of complex rearrangements. **A**) A representative intra-chromosomal complex rearrangement affecting genomic region chr15: 23.13Mb-28.62Mb in the sample S20 is shown. **B)** Inferred temporal order of events for the complex rearrangement. Associated nuclear, chromatin, replication and genomic contexts of the junctions are highlighted. All junctions have similar genomic and epigenomic contexts in this example. **C)** Proposed mechanism: Sequential DNA breaks within regions of similar replication timing, potentially triggered by a replication factory crisis. **D)** A representative intra-chromosomal complex rearrangement affecting genomic regions chr9: 62.81Mb-68.23Mb in the sample S8 is shown. **E)** Circos plot shows pairwise co-occurrences of rearrangement junctions on the same contigs. Widths of the arcs indicate optical mapping coverage. **F)** Inferred temporal order of events for the complex rearrangement. (Epi)genomic contexts of the junctions are as above.

As another case study, we examine a CGR in the in-situ tumor S8, chromosome 9, between 62.81Mb-68.23Mb that impacted the genes GXYLT1P3, TMEM252, CBWD5, FOXD4L3, FOXD4L4, LERFS, ANKRD20A3, CBWD3, and PGM5(**Figure 3D)**. There were 8 alternative DNA contigs that exhibited different combinations of structural variations, often involving common subsets of rearrangement junctions (**Supplementary Table 9**) – suggesting genetic heterogeneity. To visually represent the joint distributions of the segment junctions on the contigs, we created a circos plot, highlighting the most prevalent joint junctions: Chr9:62.81MB||Chr9:65.82MB - Chr9:65.67MB||Chr9:65.67MB, Chr9:62.81MB||Chr9:65.82MB - Chr9:63.45MB||Chr9:65.82MB, and Chr9:62.81MB|| Chr9:65.82MB - Chr9:62.90MB||Chr9:62.94MB (**Figure 3E**, **Supplementary Table 9**). Order of events analysis of this complex region indicated that the segment junctions Chr9:62.81MB||Chr9:65.82MB, Chr9:62.88MB||Chr9:62.95MB and Chr9:62.90MB||Chr9:62.94MB were the early events (**Figure 3F**), and Chr9:62.87MB||Chr9:62.96MB and Chr9_63.45MB||Chr9_65.82MB were the late events. The rearrangement junctions had different genomic and epigenomic contexts. Among the three early events, the majority are predominantly composed of strong chromatin within lamina-associated domains at the nuclear periphery. The immediate and late junctions occurred within different chromatin (strong open chromatin and weak open chromatin) and nuclear localization contexts (lamina-associated domains at the nuclear periphery and nuclear core) (**Figure 3F**). Differences between the rearrangement junctions in their allele frequencies and genomic contexts suggest that this CGR likely arose from multiple independent rearrangement events sequentially over the course of tumorigenesis. We note that genomic rearrangements may affect chromatin domain organization, which in turn may affect DNA double-strand breaks and repair pathway choices, such that the etiology of such events may be complex.

We identified 1116 CGRs in an initial genome-wide scan in our cohort; we refined the catalog based on the allele frequency of underlying rearrangements and overlap with high confidence genomic alterations as ‘major’. A total of 216 major CGRs were found in the cohort, of which 124 CGRs had all rearrangement junctions mapped. The major CGRs were uncommon in the benign lesions (0-2 per sample), but prevalent in the carcinoma in situ (0-33 per sample), locally invasive tumors (20-24), and those with distant metastasis (6-45; **Supplementary Table 10-11**). Several known cancer genes such as U2AF1, IRF4, ZNF521, PDE4DIP, HIP1, and WWTR1 were near the CGR junctions in multiple samples, however the rearrangement patterns and junctions were sample-specific. These genes also had frequent copy number alterations (2-11% cases) in the TCGA lung adenocarcinoma and squamous cell carcinoma cohorts^22^. Anyhow, based on the patterns of shared and unique rearrangement junctions, omcpIR established the order of events for 13 major CGRs. For the remaining cases, no clear order of precedence could be established reliably. It is possible that some of the latter rearrangements may have arisen due to a single event or events that were not sufficiently separated temporally, such that intermediate allelic states could be obscured by clonal sweep or low OM coverage at the junctions.

We reanalyzed the OM data using orthogonal computational frameworks to identify and annotate specific classes of complex structural variants. We used ShatterSeek^24^ to detect chromothripsis by integrating copy number and structural variation profiles. No high-confidence chromothripsis events were identified (**Supplementary Table 12**), corroborating the omcplR assessment. These observations are not unexpected, since both types of events are relatively rare. We then applied OM2BFB^25^ to the OM data to identify rearrangements stemming from breakage-fusion-bridge (BFB) cycles. The majority of detected BFB events were based on tandem duplications of a single amplicon (e.g., A, AA, AAA), which are characterized by low rearrangement junction complexity. Across the cohort, only one reached the score <1.8 to be classified as a BFB event (S20, chr2, amplicon: ABBAABBAA). We now list all putative BFB events identified in **Supplementary Table 13**. Alongside, a genome-wide scan using AmpliconReconstructorOM^26^ identified multiple potentially ecDNA-like circularized amplicon structures, all of which were single amplicon-based (**Supplementary Table 14)**. These low complexity events were limited in number and did not meet the criteria for omcplR analysis. Crucially, these results demonstrate that the high-complexity CGRs identified by omcplR represent a distinct class of genomic rearrangements that cannot be explained by established mechanisms like chromothripsis, BFB cycles, or ecDNA formation.

### (Epi)genomic contexts and DSB repair mechanisms

Analyses of the topological complexity of the order of events in the CGRs (**Figure 4A**) indicated a few common patterns. The numbers of relatively early and late events were comparable, and higher than those classified as intermediate (∼22%). This could be partially attributed to the fact that intermediate events are only present in the CGRs with order >2. Annotations of genomic contexts indicated that 73% of all junctions in CGRs had at least one of the two junction end sequences in repeat regions (**Figure 4B**). Late events were marginally enriched for both junction end-sequences in the repeat regions. While most of the junction end-sequences were either both in repeat regions (62.9%), or both in non-repetitive regions (26.9%), early events were significantly more enriched (64.5%, p-value<0.05, Binomial test; **Figure 4B**) in heterologous junctions, where one of the two junction end-sequences was in the repeat regions. Chromatin context analysis indicates that a majority of the CGR rearrangement junctions are in open chromatin, and both the junction end-sequences in open chromatin are the most common (open|open, 33.3%). Surprisingly, this was significantly more common than both the junction end-sequences in closed chromatin (closed|closed, 0.7%, p-value <0.05, Binomial test; **Figure 4B**). Taken together, repeat sequences in open chromatin may contribute towards a significant majority of rearrangement junctions in the CGRs.

**Figure 4.**
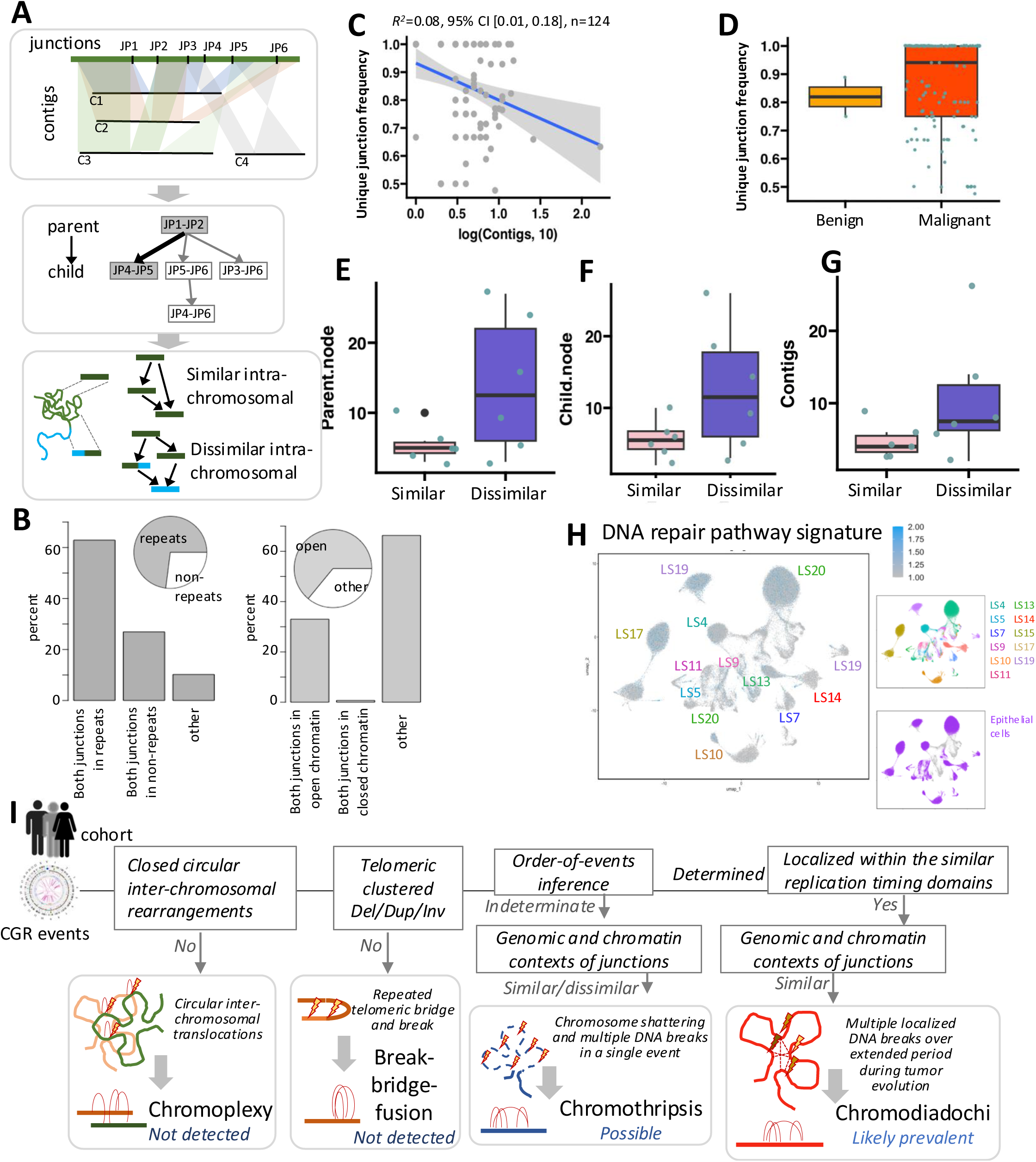
(Epi)genomic contexts and etiological inference of complex rearrangements. **A)** Schematic model illustrating the temporal ordering of rearrangement junctions in CGRs and their proposed mechanistic origins. **B**) Proportions of CGR-associated rearrangements exhibiting concordant (similar) or discordant (dissimilar) (epi)genomic contexts at junction breakpoints. **C)** Regression analysis showcasing the relationship between the number of contigs with complex structural variation junctions and the unique junctions identified. **D)** Distribution of unique complex structural variation junctions across benign and malignant (i.e. in situ, locally invasive, and distant) stages of carcinogenesis. Sequential portrayal of CGRs at the parent node (**E**), child node (**F**), and all nodes (**G**), encompassing both intra- and inter-chromosomal rearrangements alongside their chromatin annotations in Figure2D. Similar: Contigs containing identical chromatin within intra-chromosomal regions, Dissimilar: Contigs containing disparate chromatin within intra-chromosomal regions. **H**) Differences in DNA repair pathway scores in the epithelial cells from different samples based on single cell RNAseq data**. I**) Assessment of the CGRs in light of the mutagenic processes driving complex genomic alterations.

Subsequently, we investigated whether complex genomic rearrangements (CGRs) stem from heightened genomic fragility and instability within specific genomic loci. Notably, most junctions within each CGR set were unique (**Figure 4C** **and Supplementary Figure 1**). Additionally, upon stratifying by disparate cancer stages, we observed that advanced cancer stage patients exhibited a slightly higher prevalence of unique structural variation junctions, underscoring the complexity of CGRs, but the differences were not statistically significant (**Figure 4D**, Benign: 0.81**±**0.10; Malignant: 0.94**±**0.16, Mann-Whitney test p value=0.5067) and **Supplementary Figure 2**). Moreover, comparing contigs across different chromosomal levels (intrachromosomal and interchromosomal), we observed numerous CGR junctions incorporating at least one interchromosomal junction in the pre-filtered set, but this was not evident for the high-confidence set of major CGRs (**Supplementary Figure 3**; intra-chromosomal: 4.80**±**3.70, interchromosomal: 7.41**±**5.47, Mann-Whitney test p-value=3.86e-05). Further statistical analysis revealed the order of events for these junctions, indicating that parental junctions were notably complex and originated from both the same and different chromatin contexts, with a higher prevalence observed in different chromatin regions compared to identical ones (**Figure 4E**; Similar: 5.5**±**2.42; Dissimilar: 14**±**10, Mann-Whitney test p value= 0.1962; and **Supplementary Figure 4A**, one-way ANOVA p-value 2.36e-03). This trend persisted for child nodes (**Figure 4F**; Similar: 5.67**±**2.73; Dissimilar: 12.67**±**8.88, Mann-Whitney test p value= 0.1994; and **Supplementary Figure 4B**, one-way ANOVA p-value 5.54e-03) and when considering all contigs collectively (**Figure 4G**; Similar: 4.83**±**2.32; Dissimilar: 10.50**±**8.52, Mann-Whitney test p value= 0.1978) and **Supplementary Figure 4C**, one-way ANOVA p-value 0.0042), emphasizing the significance of accounting for different chromatin contexts in understanding CGR complexity.

We examined whether the clustering patterns of breakpoints within the major CGRs could be an artifact of random overlapping events or arise from non-random, localized DNA double-strand breaks due to genomic instability during tumorigenesis. For each sample, we randomly shuffled the rearrangement breakpoints within respective chromosomes preserving their chromatin, length, and genomic contexts to create simulated landscapes of genomic alterations expected by chance, repeated the simulation 10^3^ times using a published approach^27^, and each time calculated the relative frequencies of occurrence of CGRs. We found that it was significantly unlikely to observe the CGRs (simulation p-value <0.05) involving multiple (n>2) rearrangement breakpoints per sample by chance at the frequency observed in our cohort, and even more unlikely to observe the CGRs with the level of complexity observed in the in situ and advanced tumors. We observed similar results by using different strategies to create genomic rearrangements and using alternative null models in the simulation (see Methods). We further analyzed scRNAseq data for pathway-level signatures and observed that the advanced carcinomas (e.g., S17, S19, S20) with extensive CGRs also had elevated signatures of DNA repair pathways (**Figure 4H**). Although these pathway signatures are not CGR-specific, we note that active cell cycle and DNA break repair signatures in advanced carcinomas suggest continued DNA repair, which may contribute to the generation of replication-dependent or independent simple and complex rearrangements, and ultimately persistent genomic instability.

Structural inference of the CGRs allowed us to examine their likely etiologies, considering the known mutagenesis and DNA repair mechanisms broadly classified as chromoanagenesis ^6–8^. We assessed the possibilities by analyzing four attributes of the CGRs (i) intra- and inter-chromosomal CGRs, (ii) order-of-events sequence, (iii) chromosomal location, and (iv) sequence contexts of the junction regions for this purpose (**Figure 4I**). We did not find any CGR involving interconnected multiple inter- and intra-translocations and deletions, which are typical of the chromoplexy events. As noted earlier, most of the 216 major CGRs were intra-chromosomal and involved localized rearrangement events. None of those were within 5Mb of the telomere or had characteristic inversion and copy number profiles supporting break-fusion-bridge fusion (BFB). We found similar results using other methods ^25^. Among the CGRs with established order-of-events (n=13), parent and child junctions had clear allele frequency differences, which would rule out a single catastrophic event such as chromothrypsis^7,28^. A majority of these CGRs also showed branched architecture, often involving regions far away from the telomere, which makes exclusive BFB events unlikely as well. A vast majority of all rearrangement junctions in these CGRs were in low complexity or repeat sequences within gene-rich, transcriptionally active open chromatin domains, although we did not have base pair resolution information on the junction regions. Chromatin domain organization shapes replication timing^29^, such that the sequences within a chromatin domain typically replicate concurrently, and repetitive sequences permit replication-associated DNA break and repair processes^7,30^ - that is favorable for sustaining local genomic instability and fragility over multiple cell divisions during carcinogenesis; we denote it as Chromodiadochi (Greek: *chroma*: chromosome, *Diadochi*: succession). It is distinct from break-bridge-fusion events due to the absence of the characteristic inverted duplications of the amplicons, and from chromothripsis that occurs in a single shattering event^7,28^. There were other CGRs in our cohort for which we could not establish the order of events. Taken together, these data suggest that Chromodiadochimay be common. However, Chromodiadochi is not necessarily an exclusive process and can co-occur with other processes.

### Signatures of selection

Genomic alterations can gain clonal prominence by selection or drift under neutral evolution (**Figure 5A**). Allele frequency spectrum analyses indicated that a vast majority of point mutations are under neutral evolution^31^, but genomic rearrangements typically have larger effects, although a similar analysis for genomic rearrangements based on short read sequencing technology is technically challenging. The direct profiling of ultra-long DNA molecules using OM allows direct inference of allele frequency at the rearrangement breakpoints, enabling us to assess the clonal dominance and signatures of selection. Since a minor subset of the breakpoints in any CGR is early, we examined whether the CGRs are detected due to allelic abundance of early events or of the late events; in the case of the latter it would be suggestive of potential selection on the late events assuming constant mutation rate. To test that hypothesis, we first normalized the molecular support for the secondary events by the molecular support of the primary events within the respective CGR, and denoted that as the normalized allelic abundance of the late events within the CGRs. This allowed us to compare the normalized allelic abundance of the late events across samples as well as the groups of disease stages, implicitly considering the inter-chromosomal ploidy-level variations and inter-patient variations in tumor-purity and clonality of the CGRs. We found that the normalized allelic abundance of the breakpoints of the secondary events in the CGRs were significantly higher in locally advanced and distant carcinomas compared to that in the in situ tumors (**Figure 5B**; p-value <0.05, Mann-Whitney U test). Furthermore, the shape of the normalized allelic abundance distribution in the advanced tumors was skewed compared to that observed for the in-situ tumors, with an excess of secondary events with proportionally higher allelic abundance. Interpreting these observations in light of the site frequency distribution analysis, originally developed for the inference of selection on somatic point mutations^31^, we argue that the data are consistent with a relatively stronger positive selection on the secondary events in the advanced tumors.

**Figure 5:**
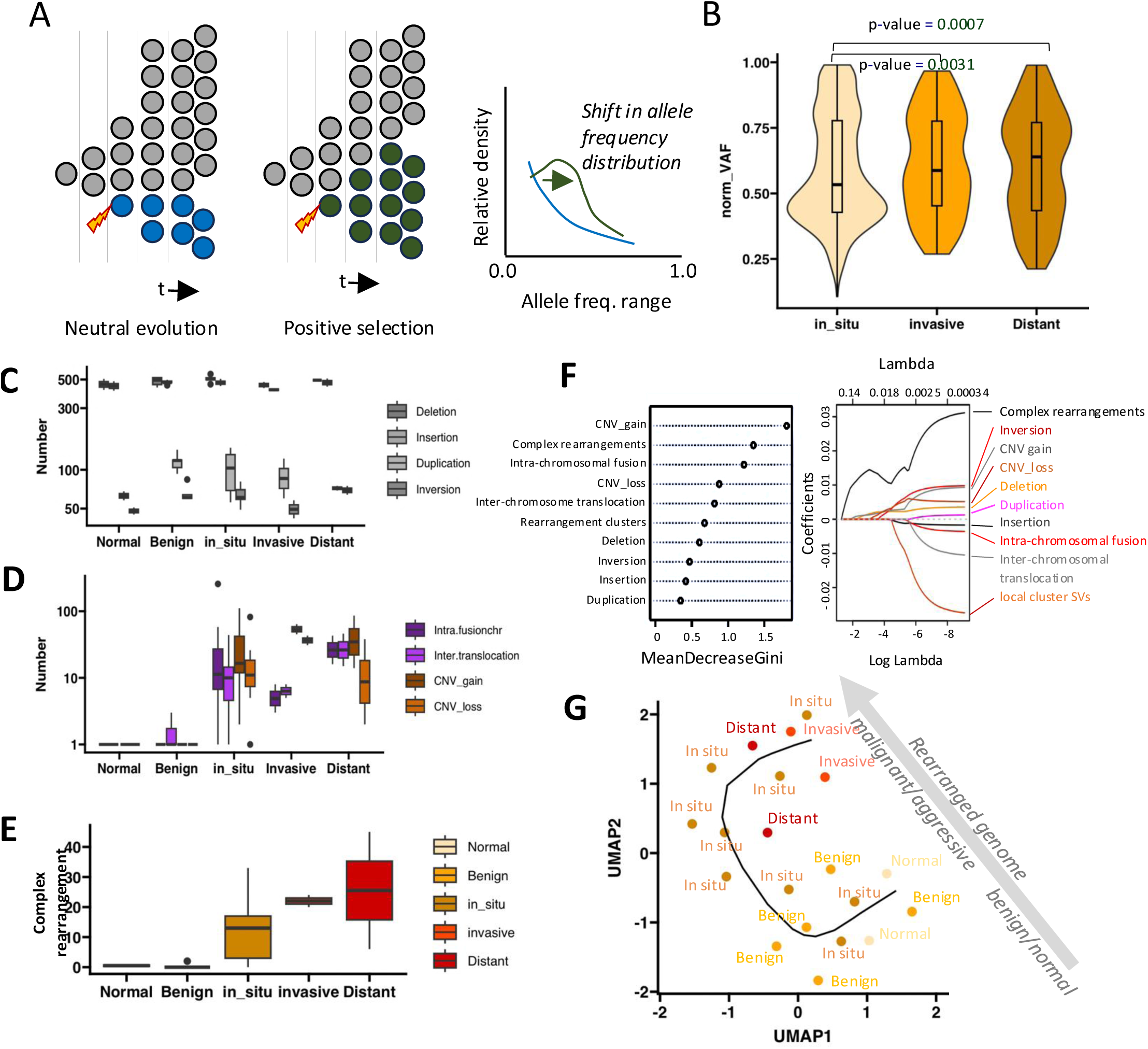
Evolutionary trajectories and progressive genomic disorganization during tumorigenesis in lung. A) A schematic representation of allele frequency change from two possible selection pressure. (B) Positive selection on cells carrying mutations result in an allele frequency distribution with an excess of high allele frequency variants. Normalized variation allele frequency for contigs observed in complex genomic rearrangements at various stages of carcinogenesis show significant increases in invasive and distant tumors. P value calculated by Kolmogorov-Smirnov test. C) Frequency of somatic genomic deletions, duplications, insertions, and inversions in the samples grouped by the stages of carcinogenesis. The frequency distribution shows modest increase with advanced disease stages. Y axis is in the log scale. D) Frequency of somatic intra-chromosomal fusions, inter-chromosomal translocations, CNV gain and losses in the samples grouped by the stages of carcinogenesis. The frequency distribution shows prevalence in normal tissues or benign lesions, but substantial increase starting with carcinoma in situ. Y axis is in the log scale. E) Frequency of somatic complex rearrangement events in the samples grouped by the stages of carcinogenesis, also shows greater prevalence with advanced disease stages. F) Random forest and Lasso feature importance for the different classes of genomic alterations in distinguishing precancerous and cancerous samples. G) Pseudotime plot showing the trajectory of decline in genome stability and organization along the clinical stages of carcinogenesis, as measured by increased burden of different classes of genomic alterations.

We integrated scRNAseq^21^ and OM data to further evaluate transcriptomic and phenotypic signatures associated with the late events in CGRs in light of known oncogenic driver genes, but the genomic resolution of the integrated dataset was typically limited to assess that conclusively. Nonetheless, several advanced tumors (e.g. S17) showed presence of transcriptionally distinct subclonal cell populations with higher proliferation scores and differences in hallmark pathway signatures indicating intra-tumor population dynamics, as reported by Venkat et al. ^21^. We note that scRNAseq data identifies functional genomic states in the tumor at the time of sample collection, while the genomic alterations in clonally amplified populations occurred much earlier during tumor development, and thus the observed associations need conservative interpretation.

### Patterns of structural variations in different disease stages

We compared the genome-wide patterns of simple and complex genomic rearrangements in the samples grouped according to their disease stages based on histopathological characteristics (**Figure 5C-E**). The burden of small structural variation events (typically 10^2^-10^4^bp; insertions, deletions, duplications, and inversions) increased in tumors compared to the non-malignant tissues, but the differences were marginal (**Figure 5C**). In contrast, copy number gain/loss and large-scale structural variations such as rearrangements within and between chromosomes were rare in precancerous tissues but orders of magnitude higher in tumors (**Figure 5D**). For instance, the normal or benign samples typically had less than 2 large copy number events, whereas advanced tumors had a notably higher number of events (e.g. 118 CNVs in S8, an in-situ tumor; **Supplementary Table 11**). Advanced tumors also exhibited a significantly higher number of intra-/inter-chromosome translocation. Notably, we detected a substantial number of 257 intra-chromosomal fusions, primarily within chromosome 8, in the in-situ sample (S14) (**Supplementary Table 11**). Somatic complex rearrangement events are more frequent in samples grouped by carcinogenesis stages, with a notable increase in prevalence in advanced disease stages (**Figure 5E**). These observations provide evidence for increased genomic disorganization at multiple levels with carcinogenesis.

### Progressive genomic disorganization with cancer stages in pseudotime

While some classes of genomic alterations were common in both precancerous and cancerous genomes, others arose predominantly in malignant stages. Lasso regression and Random Forest feature importance scores indicated that intra-/inter-chromosome translocation, copy number variations, local clustered SVs, and complex genomic rearrangements were the most significant factors that differentiated non-malignant and malignant samples. Complex genomic rearrangement had the highest coefficient in the LASSO analysis (**Figure 5F**), indicating that accumulation of complex genomic rearrangements is an important aspect of malignant disease progression.

We projected the samples, which ranged from normal lung tissues to advanced lung tumors, on the UMAP space based on the features with high coefficients in the LASSO analyses to assess the extent of variation among the samples in terms of genomic disorganization and compare that with the clinical stages of disease progression (**Figure 5G**). There were two predominant groups, one where the normal lung tissues and benign lesions were clustered together, and the other where the locally advanced and metastatic tumors were grouped together; the latter group showed greater sample-to-sample variation, indicating high inter-tumor genetic heterogeneity in advanced tumors. In situ carcinomas were the most diverse, while a subset of those were positioned with benign and normal tissues, a majority clustered with advanced malignancies, and others connected the two clusters. Thus, the UMAP projection based on the structural variation features also recapitulated the clinical stages of disease progression.

A trace of the non-linear fit through the samples represented the trajectory of progressive genomic disorganization (**Figure 5G**). Since normal tissues have mostly diploid stable genomes, and tumor genomes are highly rearranged, especially in advanced NSCLC, the path along the trajectory can be interpreted as a measure of progressive decay of genomic stability and organization in pseudotime. This analysis indicated that the velocity of decline of genomic stability varies with disease stages. While somatic genome evolution is a continuous process and many benign lesions progress to malignancies, these results suggest that the decline in genomic organization and stability may increase after malignant transformation.

## DISCUSSION

In summary, using deep-coverage single-molecule data from optical mapping (OM) technology we profiled the genomic landscapes of normal lung tissues, benign hyperplasia, dysplasia, carcinoma in situ, and advanced carcinomas. OM technology provided a major advantage, especially when compared to the standard whole genome sequencing using short read sequencing while characterizing genomic rearrangement landscapes of the tumor genomes. In general, the short read sequence technology is excellent at calling single nucleotide variants, copy number events, and small indels, which are otherwise missed by the OM technology; but it is usually insufficient for calling rearrangement junctions in repetitive regions or detecting connected genomic alteration events occurring on the same allelic sequence, especially if such events are kb-Mb apart. Indeed, we found many rearrangements using OM that were missed by short read technology, as reported elsewhere^18,19^. The long consensus maps or contigs generated by the OM technology were particularly suited for characterizing complex, intertwined genomic rearrangement events. The resolution of OM data did not allow detection of small InDels or SVs, which may accumulate in the genomic region CGR. Ultimately, a hybrid approach utilizing both short and long read technologies may help resolve the complete genome architecture and provide a reasonably complete landscape of genomic alterations in different tumor stages. Nevertheless, we found that several classes of genomic alterations, local clustered SVs and CGRs, especially inter-chromosomal complex rearrangements were uncommon in premalignant tissues but became prevalent in carcinomas. This was in stark contrast with somatic point mutations, a majority of which arise during development and aging, prior to carcinogenesis^5^.

The intricate architecture of the CGRs in these tumor genomes prompted further investigation into their underlying etiologies. We developed omcplR, a tool that utilizes genome-graph concepts to resolve CGR structures and infer their mechanistic genesis. Our analysis revealed that a subset of CGRs likely originated through progressive accumulation of individual rearrangement events in a genomic region of persistent instability. In those CGRs, the early rearrangement junctions often localized within the same chromatin domains, but the later events have more diverse contexts. We must note that the framework operates under several technical and theoretical constraints that necessitate a cautious interpretation. While computational models inherently abstract and simplify biological complexity (e.g. ploidy, rearrangement classes, 3D tumor growth, sequencing artefacts), the simulation models and Bayesian framework allowed for a systematic assessment under well-defined assumptions. We found that when the underlying events along a tumor lineage are sufficiently separated in time, the order inference is robust, but distinguishing temporally clustered events from single catastrophic events remains challenging due to potential dropouts and inherent noise in a dataset. The model inference may be sensitive to fluctuations in coverage, noise levels, or the presence of abnormal or partial contigs. The default omcplR pipeline did not consider CGR set with a small number of unique rearrangement junctions and thus may miss low complexity break-bridge-fusion events or ecDNA-like structures. Additionally, large deletions may result in a loss of information regarding prior structural states at the affected loci, potentially distorting the reconstructed evolutionary history. The network inference model relies on the Causal Markov Condition, Faithfulness, and Acyclicity. While these are standard assumptions for directed acyclic graphs in genomic modeling, their suitability should be assessed for a given dataset, especially considering biological confounders such as tumor purity, ploidy, and whole-genome doubling. OmcplR does not provide any information about the parental allelic status; it does not make any direct inference on the clonality of the CGRs and rearrangement events therein. Nonetheless, the reconstructed CGR architecture provides a structural framework to generate testable hypotheses regarding their probable molecular etiologies.

Overlaying epigenetic and replication timing data, we found interpretable associations with rearrangement junctions. Rearrangement junctions within a CGR often had similar replication timing. It is known that replication timing closely follows chromatin states^29^, and topologically associated domains tend to replicate in foci called replication factories. Replication stress in the replication factories^32–34^ is frequently associated with DNA double-strand breaks, leading us to speculate that replication factory crises may contribute to the genesis of many CGRs, especially those in the topologically associated domains with similar chromatin contexts. Evidence suggests that genomic rearrangements may drive epigenetic and chromatin changes in the proximal regions^35^, and one may argue^36^ that early genomic rearrangements may trigger downstream epigenetic changes and thereby plausibly redistribute the risk^37^ of subsequent genomic alteration events in the contexts of CGRs. We found that advanced tumors had more complex contigs and more inter-chromosomal events, which is consistent with this hypothesis, but additional work needs to be done to firmly establish the causality.

Complex rearrangements are among the least characterized classes of genomic alterations. Several mechanisms, collectively termed chromoanagenesis, have been proposed to explain these phenomena, including chromothripsis, chromoplexy, and breakage-fusion-bridge (BFB) cycles^6–8^. These processes leave distinct molecular signatures within the affected regions. In our cohort, we found no evidence of chromoplexy^7,8,38^, which typically presents as a "closed-chain" of precise inter-chromosomal translocations. Chromothrypsis^28^ is a catastrophic event involving the shattering of a chromosome followed by a random rejoining of the chromosomal fragments, typically mediated by non-homologous end-joining - resulting in the formation of clusters of complex genomic rearrangements involving intra-chromosomal translocations and deletions. Chromothripsis is usually a single event and the loci affected by chromothripsis often show genomic stability in subsequent rounds of cell divisions, such that most of the rearrangement junctions should be unique, localized and have similar ploidy-adjusted allele frequencies. In our study, the OM molecular count data at the rearrangement junctions in the CGRs suggested that many of them likely arose due to persistent genomic instability over many generations. These events were distinct from those that arise via breakage-fusion-bridge (BFB) cycles, as the identified CGRs lacked both telomeric localization and the characteristic excess of inverted duplications. Taken together, our findings using omcplR as well as other methods reveal that many CGRs arise from hierarchical combinations of localized structural variations that cannot be explained by standard chromoanagenesis models, such as break-fusion-bridge, chromothripsis, chromoanasynthesis, or chromoplexy. We propose a distinct, replication-based mechanism termed Chromodiadochi (from the Greek *Diadochi:* succession) to explain the CGRs with clearly established order-of-events in our study. Unlike the ‘one-time’ nature of chromothripsis or chromoanasynthesis, Chromodiadochi might be driven by chronic, context-dependent replication stress, that leads to recurrent DNA breaks and a cascade of nested rearrangements that accumulate over many generations during tumor evolution. While our lack of base-pair resolution precluded definitive identification of specific repair pathways, we observed that most CGR junctions were localized to repetitive sequences within transcriptionally active, open chromatin. Hence, we argue that our data is consistent with a replication-dependent model where the mutagenic process is likely driven by replication-transcription collisions that trigger fork stalling and error-prone rescue mechanisms, such as Fork Stalling and Template Switching (FoSTeS) and Microhomology-Mediated Break-Induced Replication (MMBIR) ^8,30,39^. Some CGRs with indeterminate order-of-events could potentially be due to a single mutagenesis event or multiple distinct events. We found the nested rearrangement events to be rare in premalignant tissues, and propose that Chromodiadochi may be an attribute of somatic genome evolution during malignant progression. Our observations suggest that Chromodiadochi-potentially operating in concert with other repair processes^7^ – might be a key driver of CGRs, particularly those exhibiting a clear chronological order of events, but further mechanistic studies will be needed.

The progression from premalignant lesions to carcinoma involves a fundamental shift in both cellular genomics and the coordination of the tumor microenvironment. Recent transcriptomic analyses^21^ demonstrate that alterations in immune and stromal cells not only activate oncogenic pathways but also destabilize homeostatic intercellular interactions. Our analyses of the genomic landscapes of tissues from healthy, benign and malignant lungs revealed the patterns of genome disorganization and characterized CGRs in different stages of carcinogenesis in the lung. While some classes of genomic alterations were common in both precancerous and cancerous genomes, others such as CGRs arose predominantly in malignant stages. Positive selection or even strong drift on the late events may promote the eventual clonal prominence of the CGR during tumor evolution. While tumorigenesis is a continuous process and many benign lesions progress to malignancies, these results suggest that the decline in genomic organization and instability may increase after malignant transformation. In situ tumors show substantial heterogeneity in their extent of genomic instability and disorganization in pseudotime. Given the inherent stochasticity in the accumulation of driver genetic alterations and the dynamic complexity of tumor-microenvironment interactions^40–42^, the evolutionary trajectories of such tumors may not follow a deterministic path, underscoring the clinical challenges in stratifying the patients.

## METHODS

### Cohort selection and biospecimen preparation

The cohort included pathologically normal lung tissues (*n*=2), benign pulmonary lesions (n=7), carcinoma in situ (*n*=4), locally invasive carcinoma (n=2) and metastatic lung carcinoma (*n*=3) samples (**Fig. 1A**, **Supplementary Table 1**), obtained from deidentified donors under an IRB-approved protocol. The tumors were typically resected during lobotomy or wedge resection. Tumor-adjacent pathologically normal tissues were procured from the cancer patients, while the benign pulmonary lesion samples were collected from preventive surgeries. Stains for infectious agents were negative.

### Optical genome mapping using the BioNano platform

For each sample, a minimum of 0.12g frozen tissue sample was used for the extraction and purification of ultra-high molecular weight genomic DNA (UHMW gDNA) following the manufacturer’s protocols (Bionano, San Diego, USA). Briefly, frozen tissue was homogenized in buffer, and the coarse homogenate was filtered, with the resulting suspension washed and pelleted (2200 ×g for 5 minutes), and then enzymatically treated to release gDNA. 750 ng of purified high molecular weight DNA were labeled using Direct Label Enzyme 1 (DLE-1) to conjugate DL-green fluorophores at the occurrence of a 6bp motif. After the clean-up of the excess of DL-green fluorophores and the rapid digestion of the remaining DLE-1 enzyme by proteinase K, DNA was counterstained overnight. Labeled UHMW gDNA solution (4-16ng/uL) was loaded on Saphyr Chips (Bionano, San Diego, USA) for linearization into nanochannel arrays and imaging in the Saphyr System. Images were digitized into whole-genome optical mapping (OM) molecules and label information in real time, reaching a minimum throughput target of 1500 Gbp. All samples were evaluated based on molecule size, labeling efficiency, and map rate and coverage of the GRCh38 reference.

### Structural variant detection using the rare variant analysis pipeline

Initial QC, genome alignment, and structural variation detection and copy number calling were performed using the BioNano Solve pipeline v3.7.1^16^ with default parameters. Briefly, in the automated pipeline the OM molecules of a given dataset were first aligned against GRCh38. OM molecules mapping to the same genomic regions were assembled into consensus maps. OM molecule clusters, which did not fully align with the reference diploid genome and were different from the other OM molecule clusters were identified. The Solve workflow assembled those locally into consensus maps and realigned them to the reference to generate refined and annotated SV calls. The SV calls were queried against an OGM-specific database of 179 phenotypically healthy control individuals in the pipeline to exclude potential false positives and germline variants. A given non-reference consensus map can harbor one or more SVs. It is possible to have multiple distinct consensus maps covering a region of interest, indicating different overlapping contigs carrying unique combinations of SVs. In parallel, a genome-wide copy number profile was also calculated based on depth of coverage, followed by normalization against control datasets and segmentation to identify genomic regions of copy number gain or loss, and aneusomies. For each SV and copy number variant (CNV) call, confidence scores were calculated. Variants occurring in masked regions of GRCh38 were not considered for interpretation and reporting. No inference on the parental haplotype or clonality of the SV events was drawn. Circos plots representing different classes of genomic alterations were prepared using Bionano Access v1.7.1^16^. software. Only the SVs with high allele frequency were shown in Fig. 1 for simplicity of visualization, although the downstream analyses considered all high confidence events including those present at low allele frequency.

### Short read sequencing and cross-platform comparison

Genomic DNA was extracted from 4 samples – 2 advanced lung adenocarcinoma and 2 pathologically normal tissues, and whole genome sequencing was performed using 150bp PE sequencing on an Illumina HiSeq X platform (∼30X for tumor samples, and ∼12X for the non-malignant samples, respectively). PE reads were aligned to the GRCh38Decoy build of human with BWA-MEM (v0.7.13)^43^, after removing sequencing adaptors and duplicates using fastp (v.0.21.0)^44^. Single base substitutions (SBS) and small insertions and deletions (InDels) were identified using Strelka (v2.9.2)^45^. Variants were filtered for QSS < 40(SNVs) or QSI <40 (Indels) and having at least 20× coverage in the tumor sample and 10x coverage for matched normal samples using bcftools (v1.9^46^), and at least four reads supporting the variant. The resulting variants were annotated with ANNOVAR^47^, after excluding those overlapping with centromeres and segmental duplications (genomicSuperDups) and other variants in the dbSNPv150 and 1000 Genomes Project^48^. Alongside, Manta (v1.6.0)^49^ was employed to detect structural variations (SVs) using the default parameters, followed by the extraction of the final SVs marked as ’PASS’. SV calls were compared between the BioNano OM and Illumina short read technology platforms for the 2 samples. We conducted an analysis to determine the degree of overlap between both junction ends within a 100kb range for high-confidence SVs of the same class.

### Detection of genomic regions with clusters of SVs

We filtered out the reported SVs (primarily insertions and deletions) that were shorter than 1kb. In addition, in some cases, the same SV may be represented by multiple OM molecule clusters with minor differences at the termini. To reduce such redundancy, we identified all analogous SVs that satisfy two criteria: (i) if both junction ends of the SVs fall within 5kb of one another, and (ii) the SVs are of the same type; in such cases we merged those into a single consensus SV to prepare the non-redundant list of structural variations in the somatic genomes. This practice effectively filters out technical noise in junction calling from the OM contigs as well as the minor indels or small SVs that might be at or near the major breakpoint junctions, preventing them from biasing our downstream analyses. SV clusters are ≥ 4 SVs that overlap with one another or have rearrangement junctions within 10kb distance. We analyzed the filtered set of SVs from the OM data, to identify such SV clusters, which were further examined to identify complex events. OM recorded structural variations using contigs, ensuring that each contig contains at least one structural variation. Additionally, the orientation and reference position of these structural variations are recorded. This enables us to infer the group of complex structural variations (complex rearrangement events) along with their orientation information. Prior to conducting any analysis, we performed a filtering step to exclude contigs with low confidence or potential false positives in recording structural variations. Contigs labeled as "_common" or "_segdupe", as well as those with a variant allele frequency (VAF) below 0.2, were removed. Subsequently, we retained the high-quality contigs.

### OmcplR pipeline

We developed the omcplR package to identify the complex rearrangement events in the somatic genomes. omcplr first segment the genome into equal-sized non-overlapping bins (G, default: 10kb) and track the OM contigs that contain SVs and SV junctions, which map to those bins. The genomic bins harboring multiple contigs (C, default: ≥4) are selected as initial seeds for complex rearrangement detection; typically one of those contigs represents the reference allele, while other contigs may harbor one or more genomic rearrangements. The non-reference contigs in the seed region and SV rearrangement junctions therein are retained as the initial set. Next, each contig in that set is iteratively scanned to identify additional rearrangement junctions therein and their corresponding genomic bins, which are included in the set. The process is repeated until no new non-reference contig or genomic bin is identified for a given seed, and the resulting set of contigs that harbor SVs therein denotes a complex rearrangement event. A complex event may be intra- or inter-chromosomal. In order to detect the complex rearrangement events within large structural variation events (such as copy number gain/loss and intra/inter-chromosomal structural variations) across various cancer stages, we employed the Grange function in R to overlap the initially identified complex rearrangement events with these regions. Subsequently, we tallied the retained complex rearrangement events.

A complex event with *n* contigs and *m* genomic rearrangement junctions therein can be represented as an *m* × *n* matrix to show the contig-junction relationship. Each element of the matrix is populated by the effective coverage of the OM traces of rearrangement junctions in the respective contigs. omcplR uses a Bayesian approach to determine the likely order-of-events within a complex rearrangement. It first identifies pairwise co-occurrence between the rearrangement junctions, before determining conditional relationships between the co-occurring pairs to predict potential order-of-event, and finally performing global optimization to finalize the relationship among pairs involved in a complex event using a constraint-based algorithm (PC-stable) as implemented using the bnlearn R package. The PC-stable algorithm is a constraint-based method for learning the structure of a Bayesian network. The first step is to build an undirected graph called the skeleton. This process starts with a complete undirected graph where every node is connected to every other node. Then, the algorithm systematically removes edges based on a series of conditional independence tests. Instead of immediately removing an edge as soon as an independence is found (which can affect subsequent tests for other nodes), the algorithm collects all the edges to be removed at a given step and removes them all at once, making the inferences more stable. The final step is to orient as many of the remaining undirected edges as possible without creating cycles in the graph. This process continues until no more edges can be oriented, resulting in a Completed Partially Directed Acyclic Graph (CPDAG). Each complex event is graphically represented as a Bayesian network, where the direct causal relationship junctions are connected using edges; among them, those with direct causal relationships are connected using arrows instead. Pearson’s Correlation statistic was used to calculate the significance of the connection, and a p-value < 0.05 was considered a significant difference.

To evaluate the accuracy of omcplR against known event sequences, we simulated evolutionary dynamics of clonal cell populations, modeling stochastic cell birth, death, DNA breaks, and rearrangement accumulation and catastrophic events under different parameter settings. At each time-step, the simulator preserved the rearrangement landscape of the population at single-cell resolution for lineage tracing and the order of event assessment. Simulated Complex Genomic Rearrangements (CGRs) arose via (i) sequential accumulation of genomic rearrangements along a clonal lineage, (ii) single catastrophic events, or (iii) combinations thereof. In case of a single catastrophic event, the rearrangement junctions have identical cellular counts, sequential or combined trajectories produced variable counts. A subsequent module identified clusters of overlapping rearrangements in one or more cells at a chosen time-step and reported count-level data covering the events therein and their conditional relationships – which serve as inputs for omcplr. The simulator also allowed generating alternate outputs by incorporating user-defined noise and dropout rates to vary event-level and combination-level count data, thereby enabling evaluation of model robustness. For our evaluation, we focused on loci containing two to six nested rearrangements. Each simulated complex event was analyzed either directly or after introducing varying levels of technical noise and dropouts. We then tabulated the final cellular counts and compared the inferred order of precedence against the known simulation history. Simulations were run until the simulated count data for early and late events reached levels comparable to that observed in our clinical cohort. The simulation code is available at https://github.com/sjdlabgroup/omcplr/tree/main/init to enable additional custom simulations using alternative priors, coverage profiles, and noise or dropout levels.

### Analysis of complex genomic rearrangements using different methods

We reanalyzed the OM data to examine the OM data to identify complex structural variants using established orthogonal methods. We employed ShatterSeek^24^, which allows detection of chromothripsis - a catastrophic event characterized by the shattering and random rejoining of one or more chromosomes; ShatterSeek offers unique flexibility for utilizing both copy number and structural variation calls from any genomic technology as input, and thus was suited for this analysis. We used AmpliconReconstructorOM^26^ to identify extrachromosomal DNA structures. Additionally, we ran OM2BFB^25^ with default parameters to specifically detect breakage-fusion-bridge cycles using Optical Genome Mapping data.

### Functional annotation

We annotated the genomic alterations and affected regions based on (i) genomic features––whole genes, exons, repeats, and telomeres, (ii) chromatin features – strong euchromatin, weak euchromatin, intermediate chromatin, weak heterochromatin, strong heterochromatin, (iii) nuclear localization––inter-lamina regions at the nuclear interior, and lamina-associated regions at the nuclear periphery, and (iv) replication timing based on the human reference genome hg38-using a published approach^50^. We employed the TxDb.Hsapiens.UCSC.hg38.knownGene^51^ R function to retrieve geneIDs, and used the org.Hs.eg.db package to convert the geneIDs into gene symbols.

### Analysis of transcriptomic data

We obtained processed scRNAseq data from Venkat et al.^21^ who used the 10X Chromium platform to perform single nuclei sequencing to characterize the cellular transcriptome; we note that OM and scRNAseq were performed on different tissue core specimens from the same tumors. The scRNAseq dataset was available for a subset of the samples i.e. S3-S5, S7, S9-S11, S13-S17, and S19-S20. We used this data to infer cell type composition, cellular processes, and intercellular interactions of lung tissue samples across different stages of disease progression. Pathway activity scores were calculated using approaches as described in Venkat et al.^21^.

### Evolutionary analysis and signatures of selection

To generate the expected genomic landscape of rearrangements under the null model, we shuffled the genomic rearrangements 10^3^ times preserving their chromosomes, lengths, and genomic and chromatin contexts using an approach proposed previously^27^. For inter-chromosomal translocations, we shuffled the pairs of endpoints after preserving their chromosomal assignments and genomic and chromatin contexts. We identified the CGRs using the criteria described above during simulation and analyzed their attributes to compare against the observed CGRs in the in-situ, invasive, and distant tumors. For each CGR, we designated those with the highest allele frequency (f) as the primary events (P), and others as the secondary events (S), and normalized the allele frequencies of the latter as: f’(S)=f(S)/f(P). We used the site frequency spectrum approach proposed by Williams et al.^31^ to test for neutrality for the secondary events by comparing the observed allele frequency distributions for respective CGRs against those expected under a neutral evolution model.

### Feature importance

We used LASSO (least absolute shrinkage and selection operator) a regression-based method to identify the structural variations and genomic alterations whose prevalence distinguishes between malignant and non-malignant samples in the cohort. We complemented it by using Random Forest using the randomforest R package and calculating feature importance scores.

### Evolutionary trajectory analysis

We projected the samples on the UMAP or PCA space based on selected structural variation features, which had high coefficients in the LASSO analyses. SCORPIUS^52^ was used to draw the non-linear fit on the UMAP projection of the samples. The inflection point in the plot is indicated with an arrow.

### AUTHOR INFORMATION

X.H. designed experiments, performed analyses, interpreted the results, and wrote the manuscript with input from all authors. V.V. performed single cell genomic analyses and interpreted the results. J.M., B.X., C.M., M.H., and Z.S. interpreted the results. G.R. provided pathological annotation and interpreted the results. S.D. conceived the idea, designed experiments, interpreted the results wrote the manuscript with input from all authors.

## ACKNOWLEDGEMENTS

The authors thank K.Hong, R.Mirrione, A.Biswas and J. Rosenberg for the coordination of sample collection and technical discussion.

## DATA AND CODE AVAILABILITY STATEMENT

The omcplr pipeline is available from the github repository: https://github.com/sjdlabgroup/omcplr. The genomic data are deposited to the NIH Sequence Read Archive (PRJNA574648). The source data and code are available at: https://doi.org/10.5281/zenodo.21953471. Additional processed data will be available from the corresponding author upon reasonable request..

## SUPPLEMENTARY INFORMATION

**Supplementary Figures 1-4**

**Supplementary Table 1: Sample summary for the cohort**

**Supplemental Table 2. Summary statistics of optical mapping (OM) data for the cohort**

**Supplemental Table 2. Statistics of whole genome sequencing based on short read sequencing technology**

**Supplemental Table 4. Structural variations identified using both optical mapping and whole genome sequencing**

**Supplemental Table 5. Clustered SVs identified by Bionano Optical Mapping technology**

**Supplemental Table 6. Cancer genes proximal to genomic rearrangement junctions**

**Supplemental Table 7. Assessment of the order-of-events predictions by the omcplr pipeline under different simulated conditions**

**Supplemental Table 8. Complex genomic Rearrangement (CGR) junctions in the sample S20**

**Supplemental Table 9. Complex genomic Rearrangement (CGR) junctions in the sample S8**

**Supplemental Table 10. Complex genomic Rearrangement (CGR) using different filtering criteria**

**Supplemental Table 11. Summary of different classes of structural variations**

**Supplemental Table 12. Summary of the chromothripsis events in the cohort**

**Supplemental Table 13. Summary of the break bridge fusion events in the cohort**

**Supplemental Table 14. Summary of the circular amplicons (ecDNA) in the cohort**

