## Supplementary Figure for "Deciphering the Evolutionary Histories of Complex Genomic Rearrangements in Lung Malignancies"

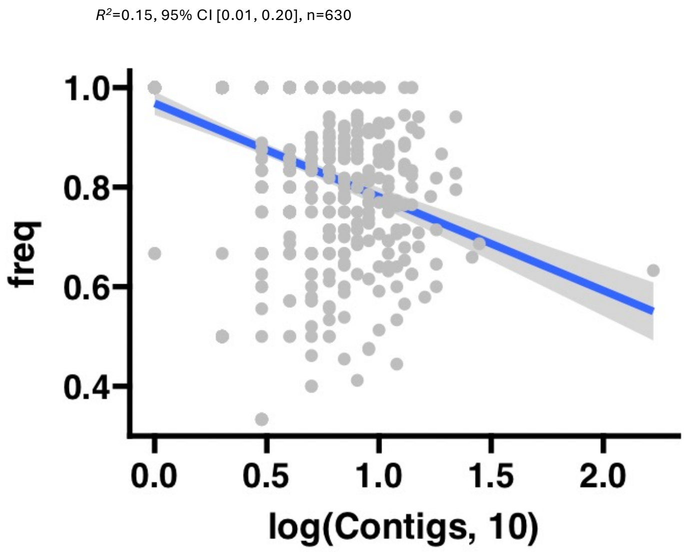


**Supplementary Figure 1.** A) Regression analysis showcasing the relationship between contigs cataloging complex structural variation junctions and the unique junctions identified.

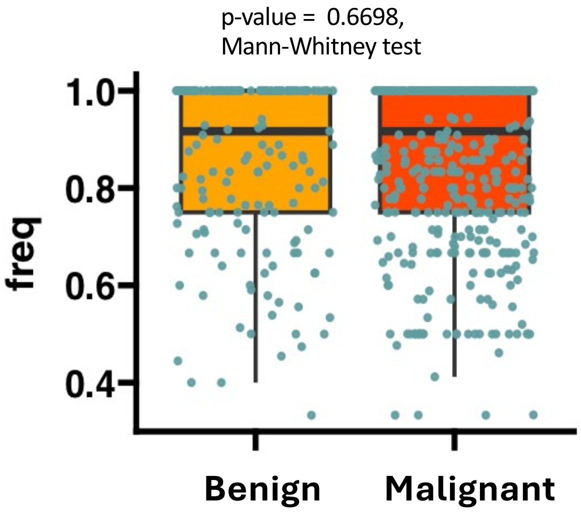


**Supplementary Figure 2.** Distribution of unique complex structural variation junctions across different stages of carcinogenesis.

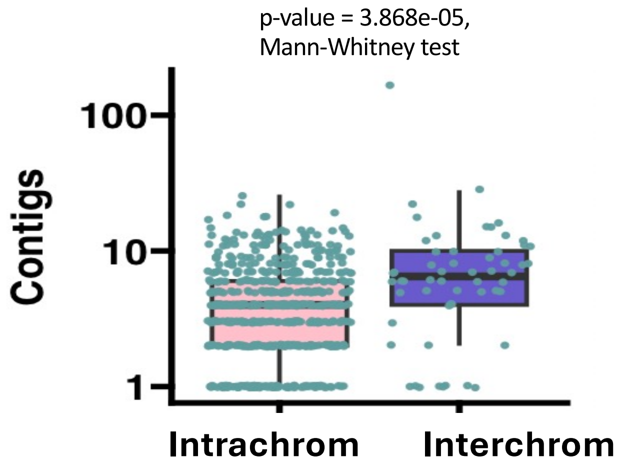


**Supplementary Figure 3.** Categorization of complex genomic rearrangement contigs based on intra- and inter-chromosomal levels.

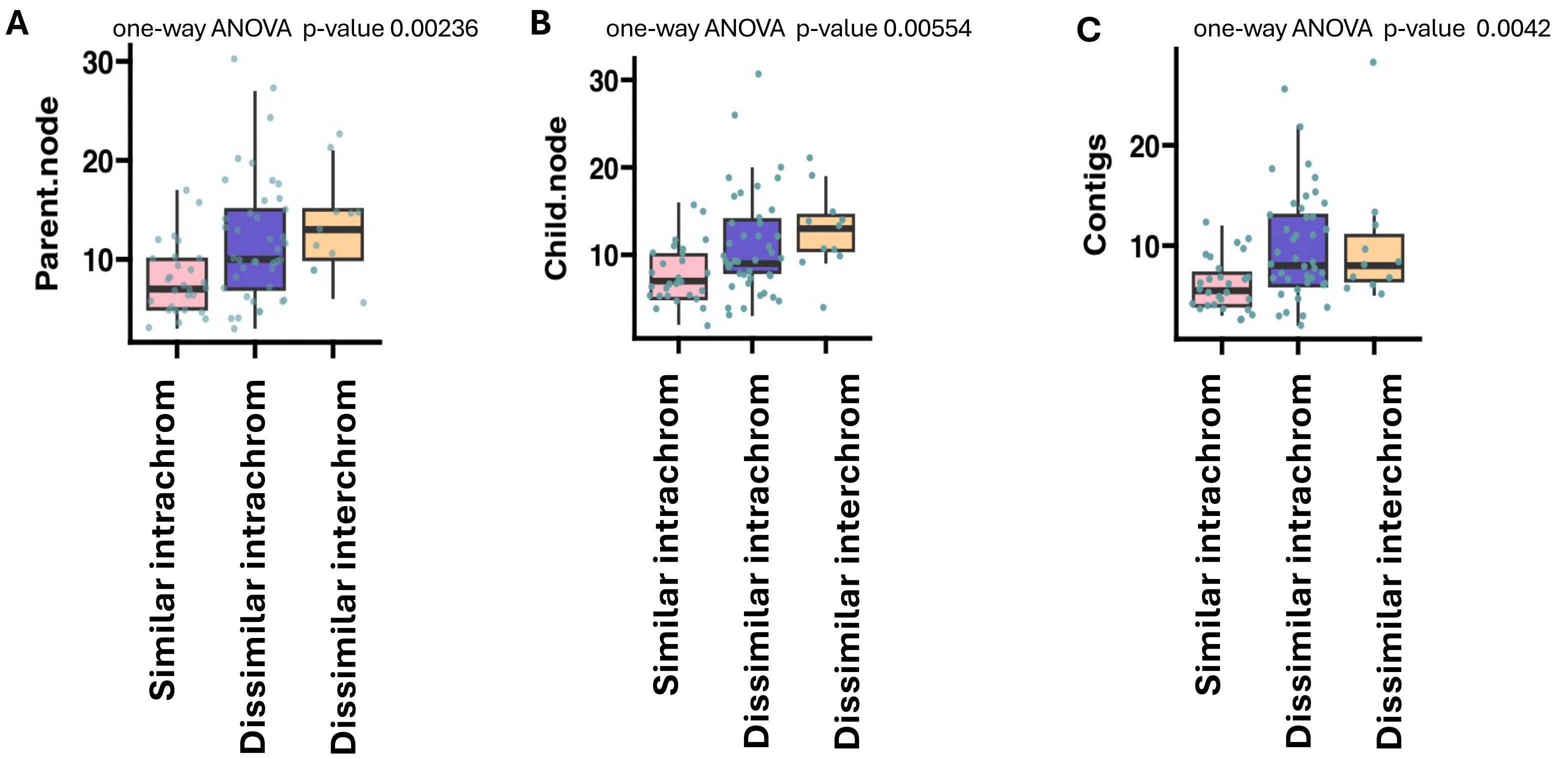


**Supplementary Figure 4.** Sequential portrayal of complex genomic rearrangement events at the parent node (A), child node (B), and all nodes (C), encompassing both intra- and inter-chromosomal rearrangements alongside their euchromatin annotations.
